# Enhanced nutrient uptake is sufficient to drive emergent cross-feeding between bacteria in a synthetic community

**DOI:** 10.1101/770727

**Authors:** Ryan K Fritts, Jordan T Bird, Megan G Behringer, Anna Lipzen, Joel Martin, Michael Lynch, James B McKinlay

## Abstract

Interactive microbial communities are ubiquitous, influencing biogeochemical cycles and host health. One widespread interaction is nutrient exchange, or cross-feeding, wherein metabolites are transferred between microbes. Some cross-fed metabolites, such as vitamins, amino acids, and ammonium (NH_4_^+^), are communally valuable and impose a cost on the producer. The mechanisms that enforce cross-feeding of communally valuable metabolites are not fully understood. Previously we engineered mutualistic cross-feeding between N_2_-fixing *Rhodopseudomonas palustris* and fermentative *Escherichia coli*. Engineered *R. palustris* excreted essential nitrogen as NH_4_^+^ to *E. coli* while *E. coli* excreted essential carbon as fermentation products to *R. palustris*. Here, we enriched for nascent cross-feeding in cocultures with wild-type *R. palustris*, not known to excrete NH_4_^+^. Emergent NH_4_^+^ cross-feeding was driven by adaptation of *E. coli* alone. A missense mutation in *E. coli* NtrC, a regulator of nitrogen scavenging, resulted in constitutive activation of an NH_4_^+^ transporter. This activity likely allowed *E. coli* to subsist on the small amount of leaked NH_4_^+^ and better reciprocate through elevated excretion of organic acids from a larger *E. coli* population. Our results indicate that enhanced nutrient uptake by recipients, rather than increased excretion by producers, is an underappreciated yet possibly prevalent mechanism by which cross-feeding can emerge.

## INTRODUCTION

Microorganisms typically exist as members of diverse and interactive communities wherein nutrient exchange, also known as cross-feeding, is thought to be ubiquitous [1-7]. The prevalence of cross-feeding might explain, in part, why many microbes cannot synthesize essential vitamins and amino acids (i.e. auxotrophy), as they can often acquire these compounds from other community members [1, 7, 8]. Furthermore, microbes in nature experience varying degrees of starvation and often exist in states of low metabolic activity [9, 10]. Thus, cross-feeding might also serve to sustain microbes through starvation. Despite the prevalence of cross-feeding, elucidating the molecular mechanisms underlying emergent cross-feeding interactions and tracking their evolutionary dynamics within natural microbial communities is difficult due to their sheer complexity. To overcome this intrinsic complexity, tractable synthetic consortia have proven useful for studying aspects of the mechanisms, ecology, evolution, and applications of microbial communities [4, 11-16].

To study the molecular mechanisms of nutrient cross-feeding, we previously developed a bacterial coculture in which *Escherichia coli* and *Rhodopseudomonas palustris* reciprocally exchange essential metabolites under anaerobic conditions (Fig.1A) [17-20]. In this coculture, *E. coli* ferments glucose, a carbon source that *R. palustris* cannot consume, and excretes ethanol and organic acids, namely acetate, lactate, succinate, and formate, as waste products. The organic acids, with the exception of formate, serve as the sole carbon sources for *R. palustris* (Fig. 1A). In return, *R. palustris* fixes dinitrogen gas (N_2_) via the enzyme nitrogenase and excretes ammonium (NH_4_^+^), which is the sole nitrogen source for *E. coli* (Fig. 1A). Because both species depend on essential nutrients provided by their partner, this coculture functions as a synthetic obligate mutualism.

**Fig. 1.**
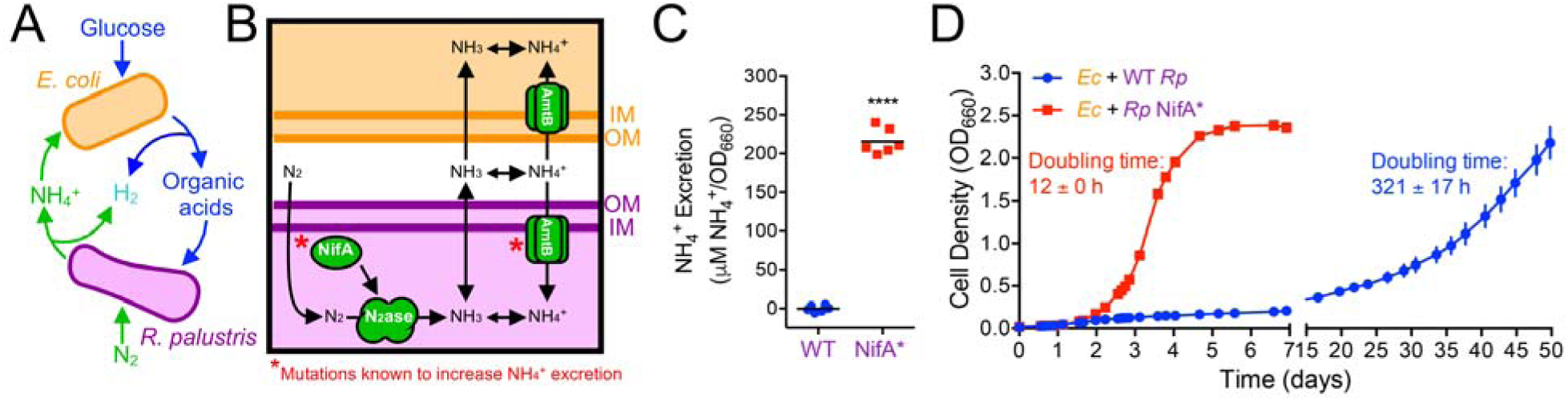
Mutualistic cross-feeding between *E. coli* and *R. palustris* is facilitated by NH_4_^+^ excretion. (A) Coculture growth requires reciprocal cross-feeding of organic acids and NH_4_^+^ excreted by *E. coli* and *R. palustris*, respectively. (B) Mechanism of NH_4_^+^ cross-feeding from *R. palustris* to *E. coli* and mutational targets known to increase NH_4_^+^ excretion by *R. palustris* (*). (C) NH_4_^+^ excretion levels by WT *R. palustris* (CGA009) and an isogenic NifA* mutant (CGA676) in carbon-limited N_2_-fixing monocultures grown in grown in MDC or NFM minimal medium, with similar results observed for both media. Points are biological replicates and lines are means, n=6; paired t-test, \*\*\*\**p*<0.0001. (D) Coculture growth curves (both species) of *E. coli* paired with either WT *R. palustris* or the NifA* mutant. Points are means ± SEM, n=3. Doubling times are means ± SD.

NH_4_^+^ cross-feeding from *R. palustris* to *E. coli* is thought to depend on the equilibrium between NH_3_ and NH_4_^+^. The small proportion of NH_3_ present in neutral pH environments is membrane permeable and can diffuse out of cells [21, 22]. Leaked NH_4_^+^ can be recaptured by AmtB transporters [21], which in the case of *R. palustris* helps privatize valuable NH_4_^+^ (Fig. 1B) [17, 18]. NH_4_^+^ leakage is also limited through the strict regulation of N_2_ fixation, including by the transcriptional activator NifA, so that energetically expensive N_2_ fixation is only performed when preferred nitrogen sources, such as NH_4_^+^, are limiting [23]. Previously, we identified two types of mutations that increase NH_4_^+^ excretion by *R. palustris* during N_2_ fixation and support coculture growth with *E. coli* [17]: (i) deletion of *amtB*, which prevents recapture of leaked NH_3_, or (ii) a 48-bp deletion within *nifA* (denoted as NifA*), which locks NifA into an active conformation [24] (Fig. 1B). In contrast, wildtype *R. palustris* does not readily support coculture growth with *E. coli* due to insufficient NH_4_^+^ excretion [17].

While mutualistic cross-feeding of communally valuable NH_4_^+^ between *E. coli* and *R. palustris* can be rationally engineered, we questioned whether such an interaction could arise spontaneously. Herein we experimentally evolved cocultures pairing WT *E. coli* with either WT *R. palustris* or an engineered NifA* mutant in serially-transferred batch cultures for ∼150 generations. In both cocultures, a mutualism was established and growth rates improved over serial transfers, but growth and metabolic trends remained distinct. By pairing ancestral and evolved isolates of each species, we determined that adaptation by *E. coli* was solely responsible for establishing a mutualism with WT *R. palustris*. Whole-genome sequencing and subsequent genetic verification identified a missense mutation in the *E. coli* transcriptional activator for nitrogen scavenging, NtrC, that was sufficient to enforce mutualistic NH_4_^+^ cross-feeding with WT *R. palustris*. This mutation results in constitutive AmtB expression, presumably enhancing NH_4_^+^ uptake. Our results suggest that mutations that improve acquisition of communally valuable nutrients by recipients are favored to evolve and can promote the emergence of stable cross-feeding within synthetic consortia, and potentially within natural communities.

## MATERIAL AND METHODS

### Bacterial strains and growth conditions

All strains and plasmids are listed in Supplementary Table S1. All *E. coli* strains used in this study are derived from the type strain MG1655 [25], unless noted otherwise. The WT and NifA* *R. palustris* strains used in Fig. 1 were the type strain CGA009 [26] and CGA676, respectively. CGA676 carries a 48 bp deletion in *nifA* [24]. The *R. palustris* strains used in experimental coculture evolution and subsequent experiments were CGA4001 and CGA4003, which are derived from CGA009 and CGA676, respectively, with both carrying an additional Δ*hupS* mutation, preventing H_2_ oxidation.

*E. coli* was grown in lysogeny broth (LB)-Miller (BD Difco) or on LB plates with 1.5% agar at 30 or 37°C with gentamicin (Gm; 5-15 µg/ml), kanamycin (Km; 30 µg/ml), or carbenicillin (Cb; 100 µg/ml) when appropriate. *R. palustris* was grown in defined minimal photosynthetic medium (PM) [26] or on PM agar with 10 mM succinate at 30°C with Gm (100 µg/ml) when appropriate. N_2_-fixing medium (NFM) was made by omitting (NH_4_)_2_SO_4_ from PM. NFM and LB agar were used as selective media to quantify *R. palustris* and *E. coli* colony-forming units (CFUs), respectively. Experimental mono- and cocultures were grown in 10 ml of M9-derived coculture medium (MDC) in 27 ml anaerobic glass test tubes. Tubes were made anaerobic under 100% N_2_, sterilized, and supplemented with 1 mM MgSO_4_ and 0.1 mM CaCl_2_ as described [17]. *E. coli* starter monocultures had 25 mM glucose and were growth-limited by supplementing with 1.5 mM NH_4_Cl. *R. palustris* starter monocultures were growth-limited by supplementing with 3 mM acetate. Cocultures were inoculated by subculturing 1% v/v of starter monocultures of each species into MDC with 50 mM glucose. Mono- and cocultures were grown at 30 °C, under shaken conditions, lying horizontally and shaken at 150 rpm beneath a 60 W incandescent bulb (750 lumens), or under static conditions, standing vertically without shaking beside a 60 W incandescent bulb.

### *E. coli* strain construction

All primers are listed in Supplementary Table S2. To construct the *E. coli* NtrC^S163R^ mutant, the Gm^R^-*sacB* genes from pJQ200SK [27] were PCR amplified using primers containing ∼40 bp overhangs with homology up- and downstream of *ntrC* (*glnG*). A second round of PCR was subsequently performed to increase the length of overhanging regions of homology to ∼80 bp to increase the recombination frequency. *E. coli* harboring pKD46, encoding arabinose-inducible λ-red recombineering genes [28], was grown in LB with 20 mM arabinose and Cb at 30 °C to an OD_600_ of ∼0.5 and then centrifuged, washed, and resuspended in sterile distilled water at ambient temperature. Resuspended cells were electroporated with the Gm^R^-*sacB* PCR product containing overhangs flanking *ntrC* and plated on LB Gm agar. Gm-resistant colonies were screened by PCR for site-directed recombination of Gm^R^-*sacB* into the *ntrC* locus, creating a Δ*ntrC*::Gm^R^-*sacB* allele, which was then verified by sequencing. To replace the Δ*ntrC*::Gm^R^-*sacB* locus, the NtrC^S163R^ allele was PCR-amplified from gDNA from evolved *E. coli* (lineage A25) and electroporated into *E. coli* Δ*ntrC*::Gm^R^-*sacB* harboring pKD46. After counterselection on LB agar with 10% (w/v) sucrose but without NaCl, site-directed recombination of the NtrC^S163R^ allele into the native locus was confirmed by PCR and sequencing. *E. coli* NtrC^S163R^ was grown overnight on LB agar at 42°C to cure the strain of pKD46, which was confirmed by Cb sensitivity.

### *R. palustris* strain construction

To construct *R. palustris* CGA4001 and CGA4003, pJQ-Δ*hupS* was introduced into *R. palustris* CGA009 and CGA676, respectively, by conjugation with *E. coli* S17-1. Mutants were then obtained using sequential selection and screening as described [29]. The Δ*hupS* deletion was confirmed by PCR and sequencing.

### Analytical procedures

Cell densities were approximated by optical density at 660 nm (OD_660_) using a Genesys 20 visible spectrophotometer (Thermo-Fisher). Coculture doubling times were derived from specific growth rates determined by fitting exponential functions to OD_660_ measurements between 0.1-1.0 for each biological replicate. NH_4_^+^ was quantified using an indophenol colorimetric assay [17]. Glucose and soluble fermentation products were quantified by high-performance liquid chromatography (Shimadzu) as described [30]. H_2_ was quantified by gas chromatography (Shimadzu) as described [31].

### Coculture evolution experiments

Founder monocultures of *E. coli* MG1655, *R. palustris* CGA4001 (Δ*hupS*), and CGA4003 (Δ*hupS* NifA*) were inoculated from single colonies in MDC. Once grown, a single founder monoculture of each strain was used to inoculate twelve WT-based cocultures (six shaken: A-F; six static: G-L) and 12 NifA*-based cocultures (six shaken: M-R; six static: S-X) in MDC with 50 mM glucose. Cocultures were serially transferred by passaging 2% v/v of stationary phase coculture (OD_660_ > 2 and a low metabolic rate based on H_2_ measurements) into fresh MDC. The NifA*-based cocultures were transferred weekly whereas WT-based cocultures were transferred every 21-50 days for the first five transfers and then every two-weeks based on the time required to reach OD_660_ > 2. For comparative analyses, shaken cocultures (A-F and M-R) were revived from frozen stocks made following transfer-2 (generation 17) and transfer-25 (generation 146). Frozen stock (∼0.2 ml) was thawed in 1 ml sterile MDC, washed 2X with MDC to remove glycerol, and then resuspended in 0.2 ml MDC for use as inoculum.

### RNA extraction and reverse transcription quantitative PCR

RNA was isolated from exponentially growing *E. coli* monocultures or starved cell suspensions that had been chilled on ice, centrifuged at 4°C, cell pellets frozen using dry-ice in ethanol, and stored at -80°C. Cell pellets were thawed on ice, disrupted by bead beating, and then RNA was purified using an RNeasy MiniKit (Qiagen), Turbo DNase (Ambion) treatment on columns, and RNeasy MinElute Cleanup Kit (Qiagen). cDNA was synthesized from 0.5-1 µg of RNA per sample using Protoscript II RT and Random Primer Mix (New England Biolabs). qPCR reactions were performed on cDNA using iQ SYBR Green supermix (BioRad). *E. coli* gDNA was used to generate standard curves for *amtB* and *ntrC* transcript quantification, which were normalized to transcript levels of reference genes *gyrB* and *hcaT* [32]. Two technical replicate qPCR reactions were performed and averaged for each biological replicate to calculate relative expression.

### Genome sequencing and mutation analysis

gDNA was extracted from stationary phase evolved cocultures following revival from frozen stocks using a Wizard Genomic DNA purification Kit (Promega). DNA fragment libraries were constructed for samples from shaking WT-based cocultures A-F and NifA*-based cocultures M-R at generation ∼146 using NextFlex Bioo Rapid DNA kit. Samples were sequenced on an Illumina NextSeq 500 150 bp paired-end run by the Indiana University Center for Genomics and Bioinformatics. Paired-end reads were trimmed using Trimmomatic 0.36 [33] with the following options: LEADING:3 TRAILING:3 SLIDINGWINDOW:10:26 HEADCROP:10 MINLEN:36. Mutations were called using *breseq* version 0.32.0 on Polymorphism Mode [34] and compared to a reference genome created by concatenating *E. coli* MG1655 (Accession NC_000913), *R. palustris* CGA009 (Accession BX571963), and its plasmid pRPA (Accession BX571964). Mutations are summarized in Supplemental File 1.

Additional gDNA sequencing for evolved WT-based cocultures A-F (shaking, generation 11), G-L (static, generations 11 and 123), and NifA*-based cocultures S-X (static, generation 123) was performed at the US Department of Energy Joint Genome Institute. Plate-based DNA library preparation for Illumina sequencing was performed on the PerkinElmer Sciclone NGS robotic liquid handling system using Kapa Biosystems library preparation kit. 200 ng of gDNA was sheared using a Covaris LE220 focused-ultrasonicator. Sheared DNA fragments were size selected by double-SPRI and selected fragments were end-repaired, A-tailed, and ligated with Illumina compatible sequencing adaptors from IDT containing a unique molecular index barcode for each sample library. Libraries were quantified using KAPA Biosystem’s next-generation sequencing library qPCR kit and run on a Roche LightCycler 480 real-time PCR instrument. The quantified libraries were then prepared for sequencing on the Illumina HiSeq sequencing platform utilizing a TruSeq Rapid paired-end cluster kit. Sequencing was performed on the Illumina HiSeq2500 sequencer using HiSeq TruSeq SBS sequencing kits, following a 2×100 indexed run recipe. Reads were aligned to a reference genome created by concatenating *E. coli* MG1655 (Accession NC_000913), *R. palustris* CGA009 (Accession NC_005296), and its plasmid pRPA (Accession NC_005297) [35]. The resulting bams were then split by organism and down sampled to 100 fold depth if in excess of that, then re-merged to create a normalized bam for calling single nucleotide polymorphisms and small indels by callVariants.sh from the BBMap package (sourceforge.net/projects/bbmap/) to capture variants present within the population and annotation applied with snpEff [36]. Mutations are summarized in Supplemental File 2.

All FASTQ files are available at NCBI Sequence Read Archive (Accession numbers listed in Supplementary Table S3)

## RESULTS

### Nascent emergence of mutualistic cross-feeding between wild-type *R. palustris* and *E. coli*

Previously, we engineered *R. palustris* to excrete NH_4_^+^ (NifA*) to stabilize mutualistic cross-feeding with *E. coli* (Fig. 1A and B). Here, we sought to determine whether such a relationship could evolve spontaneously. Spatial proximity has been shown to be an important factor in many microbial cross-feeding mutualisms [37, 38], which can be disrupted by mixing. To account for the possible importance of proximity, we established cocultures with WT *R. palustris* (WT-based cocultures) under both shaken conditions, wherein cells are evenly distributed, and static conditions, wherein cells settle in close proximity at the bottom of the tube. In parallel, we also established shaken and static cocultures featuring the *R. palustris* NifA* strain (NifA*-based cocultures) as a comparative reference.

We confirmed our previous observations [17] that WT *R. palustris* exhibits undetectable NH_4_^+^ excretion and does not readily support coculture growth with WT *E. coli*, in contrast to the NifA* mutant (Fig. 1C and D). Whereas shaken NifA*-based cocultures grew to an OD_660_ > 2.0 in 4-6 days with a doubling time of ∼12 h, shaken WT-based cocultures did not exhibit appreciable growth in the same time frame (Fig. 1D). We hypothesized that prolonged incubation might enrich for spontaneous mutants that permit coculture growth. Indeed, after 50 days, shaken WT-based cocultures reached densities similar to those observed for NifA*-based cocultures, albeit with a doubling time of ∼13 days (Fig. 1D). Static WT- and NifA*-based cocultures also became turbid within this time frame.

Upon observing a nascent mutualism between WT *R. palustris* and *E. coli*, we set up new cocultures to experimentally evolve six replicates of WT-based cocultures (A-F) and NifA*-based cocultures (M-R) through serial transfers under shaken conditions, all with WT *E. coli* (Fig. 2A), to compare their physiology, evolutionary trajectory, and species and genotypic composition. We also serially transferred static WT-based cocultures (G-L) and NifA*-based cocultures (S-X). However, we herein focus the bulk of our analyses on shaken cocultures because (i) the close proximity provided under static conditions was not required for nascent cross-feeding, (ii) shaken conditions facilitate analyses such as OD-based determination of growth rates, and (iii) strong biofilms emerged in static cocultures, complicating the determination of population densities.

**Fig. 2.**
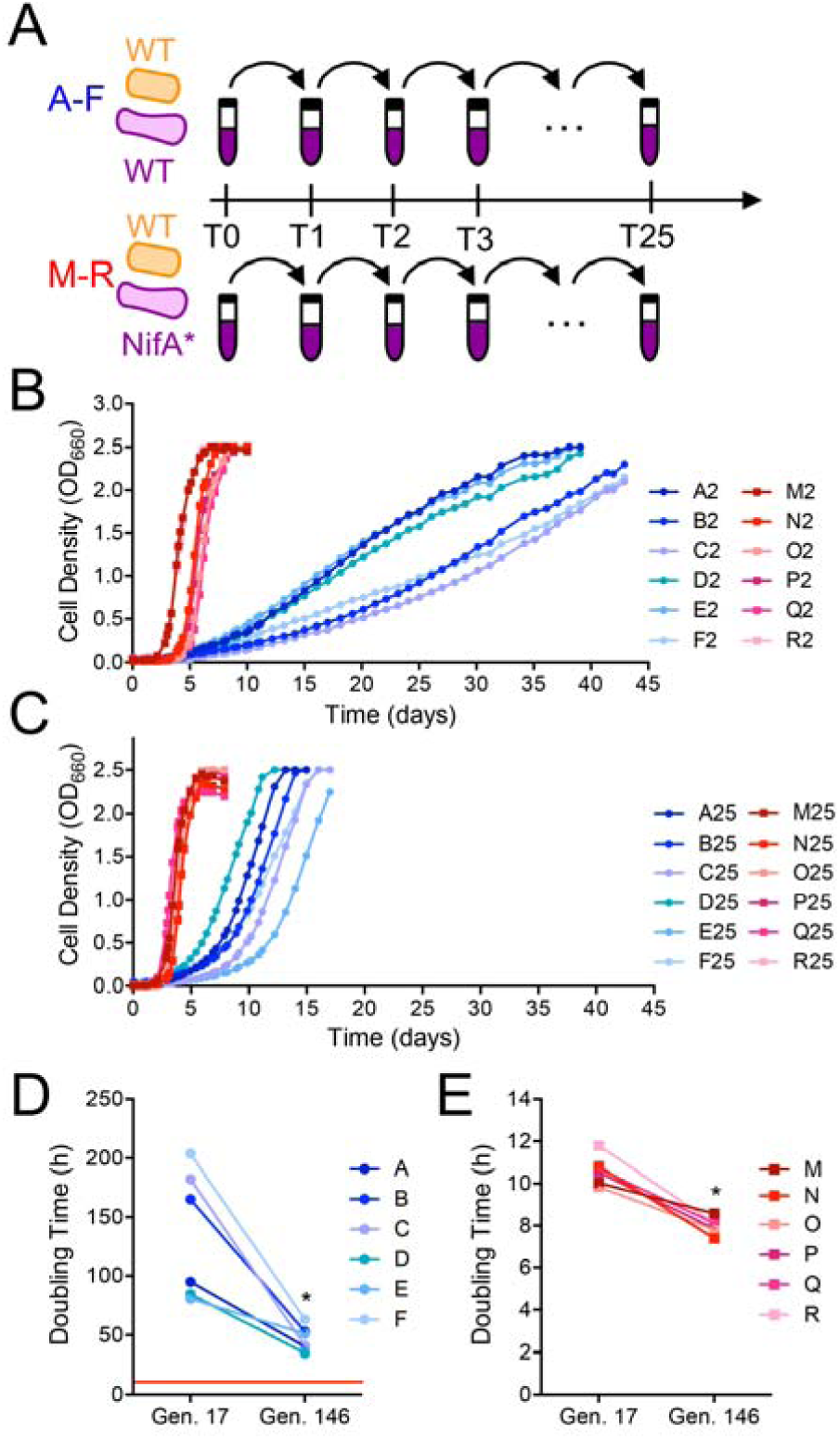
Coculture doubling times decreased during experimental evolution of WT-based (CGA4001; blue) and NifA*-based (CGA4003; red) cocultures. Points are values for the indicated individual revived coculture lineages. (A) Design for experimental evolution of parallel WT-based (A-F) and NifA*-based (M-R) cocultures via serial transfer. (B, C) Growth curves (both species) of WT-based (blue circles) and NifA*-based (red squares) cocultures revived after two transfers (17 generations) (B) or 25 transfers (146 generations) (C) of experimental evolution. Different shades indicate the different lineages. (D, E) Coculture doubling times (both species) of individual WT-based cocultures (D) or NifA* based cocultures (E) at generation (Gen.) 17 and 146 (*, Wilcoxon matched-pairs signed rank, *p*=0.0313). (D) The red line indicates the doubling time of NifA*-based cocultures at Gen. 17.

Shaken WT- and NifA*-based cocultures were serially transferred 25 times, corresponding to ∼146 generations, with ∼5.6 generations estimated per serial coculture (including the original cocultures designated, transfer-0) based on the 1:50 dilution used for each transfer. This number of generations corresponded to ∼ 65 weeks for WT-based cocultures and ∼ 26 weeks for NifA*-based cocultures. We then revived cocultures from frozen stocks at transfer-2 (generation 17; G17) and transfer-25 (generation 146; G146) time point to compare growth and population trends. At G17, NifA*-based cocultures exceeded an OD_660_ of 2 in under 8 days whereas WT-based cocultures took ∼40 days (Fig. 2B). By G146, the time needed to reach OD_660_ > 2 had decreased for every lineage (Fig. 2C). The shortened growth phase was most pronounced for WT-based cocultures, which all reached OD_660_ > 2 in under 17 days by G146, less than half the time needed at G17 (Fig. 2B, C); WT-based coculture doubling times decreased from 135 ± 55 h to 47 ± 10 h (Fig. 2D). Though less drastic, NifA*-based coculture doubling times also decreased, in this case from ∼11 h to ∼8 h (Fig. 2D). Thus, WT-based cocultures adapted to grow faster, although never as fast as unevolved engineered NifA*-based cocultures.

Because growth trends differed between WT- and NifA*-based cocultures, we wondered how species populations were affected. We therefore enumerated viable cells as colony forming units (CFUs) at the final time points for G17 and G146 cocultures shown in Fig. 2A and B. At G17, both *R. palustris* and *E. coli* populations in WT-based cocultures were lower than those in NifA*-based cocultures (Fig. 3A). It is worth noting that NifA*-based cocultures were plated after ∼10 days, whereas WT-based cocultures were plated after 39-43 days due to their slower growth rate. Consequently, the background death rate during the additional ∼30 days of slower growth for WT-based cocultures could have contributed to the lower final CFUs. At G146, *R. palustris* abundances in WT-based cocultures had increased >14-fold and exceeded *R. palustris* abundances observed in NifA*-based cocultures by ∼2-fold (Fig. 3). *E. coli* abundances in WT-based cocultures also increased >7-fold by G146, but did not reach abundances observed in NifA*-based cocultures (Fig. 3). The increase in *E. coli* abundances in WT-based cocultures by G146 suggests that *E. coli* had better access to NH_4_^+^, or other nitrogen compounds than at G17 (Fig. 3B). Due to the disproportionate increase in each population in WT-based cocultures between G17 and G146, *E. coli* percentages remained low at 1-5%, relative to 11-21% in NifA*-based cocultures (Fig. 3B). These differences in *E. coli* populations between WT- and NifA*-based cocultures are consistent with previous findings that higher NH_4_^+^ excretion by *R. palustris* supports faster growth and higher *E. coli* abundances [17-19].

**Fig. 3.**
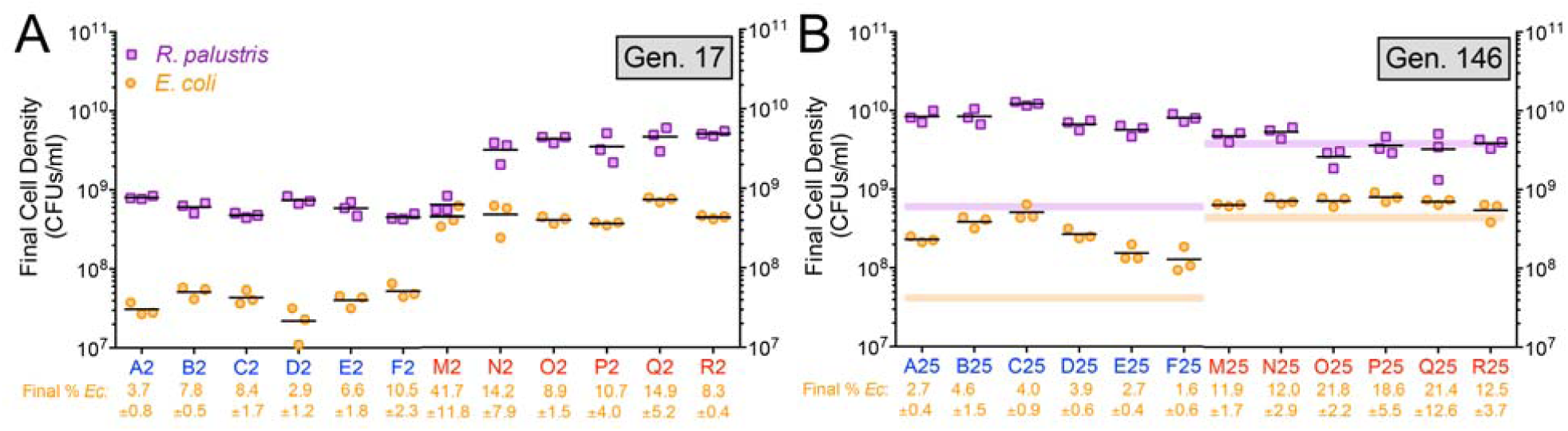
Final cell densities for each species increase in WT-based cocultures between generation 17 (A) and generation 146 (B). (A, B) Final viable cell densities (colony forming units [CFUs/ml]) of *R. palustris* and *E. coli* and the final *E. coli* percentage (± SD) for WT-based (CGA4001; blue) and NifA*-based (CGA4003; red) cocultures at the final time points shown in Fig 2B and 2C. Points represent technical replicates of CFUs/ml for each lineage and lines are means, n=3. The purple and orange lines in panel B indicate the median CFUs/ml for *R. palustris* and *E. coli*, respectively, at generation 17 for reference. The lower *R. palustris* CFUs in the M2 coculture was due to plate contamination that obscured accurate CFU enumeration. The average final *E. coli* percentage did not differ significantly between WT-based and NifA*-based cocultures at Gen. 17 whether we included or excluded M2 values (Wilcoxon matched-pairs signed rank test, *P*=0.563 or 0.125).

In contrast to WT-based cocultures, NifA*-based cocultures did not display drastically higher cell densities for each species between G17 and G146 (Fig. 3). Average *E. coli* densities were 4.7 x 10^8^ CFUs/ml and 6.9 x 10^8^ CFUs/ml at G17 and G146, respectively. Average *R. palustris* densities were 3.4 x 10^9^ CFUs/ml and 3.9 x 10^9^ CFUs/ml at G17 and G146, respectively. The average *E. coli* percentage in NifA*-based cocultures was similar at G17 (16.5%) and at G146 (16.4%) (Wilcoxon matched-pairs signed rank test, *P*=0.563).

### Metabolic differences between WT- and NifA*-based cocultures help explain growth and population trends

Our studies demonstrated that growth and population trends in coculture are strongly influenced by cross-feeding levels of both NH_4_^+^ and organic acids [17-19]. For instance, higher NH_4_^+^ cross-feeding to *E. coli* leads to higher *E. coli* growth rates [17-19]. In turn, higher *E. coli* growth rates boost organic acid excretion, supporting better *R. palustris* growth up until the organic acid excretion rate exceeds the rate at which *R. palustris* can consume them [17]. To see if such trends were also present in WT-based cocultures, we quantified glucose consumption and fermentation product yields at G17 and G146 at stationary phase. At G17, glucose consumption by *E. coli* in WT-based cocultures was about half of that in NifA*-based cocultures (Fig. 4A). The lower glucose consumption in WT-based cocultures can explain in part the lower *E. coli* CFUs observed in Fig. 3A. Previously, we showed that non-growing *E. coli* can ferment glucose, and that this growth-independent fermentation can provide sufficient carbon to support *R. palustris* growth [19]. We hypothesize that growth-independent fermentation by *E. coli* was an important cross-feeding mechanism during the extremely slow growth of early WT-based cocultures (Fig. 1D). However, by G146, *E. coli* glucose consumption in WT-based cocultures approached that in NifA*-based cocultures in most lineages, with one lineage consuming more glucose than NifA*-based cocultures (Fig. 4A). The general increase in *E. coli* glucose consumption in WT-based cocultures from G17 to G146 likely supported both the faster coculture doubling times (Fig. 2D) and higher *E. coli* abundances (Fig. 3B) and contributed to the accumulation of consumable organic acids such as acetate and succinate in some WT-based cocultures at G146, qualitatively similar to NifA*-based cocultures (Fig. S2). Overall, the increase in metabolic activities attributed to *E. coli* and the improved *E. coli* growth in WT-based cocultures between G17 and G146 suggests that nitrogen cross-feeding also increased.

**Fig. 4.**
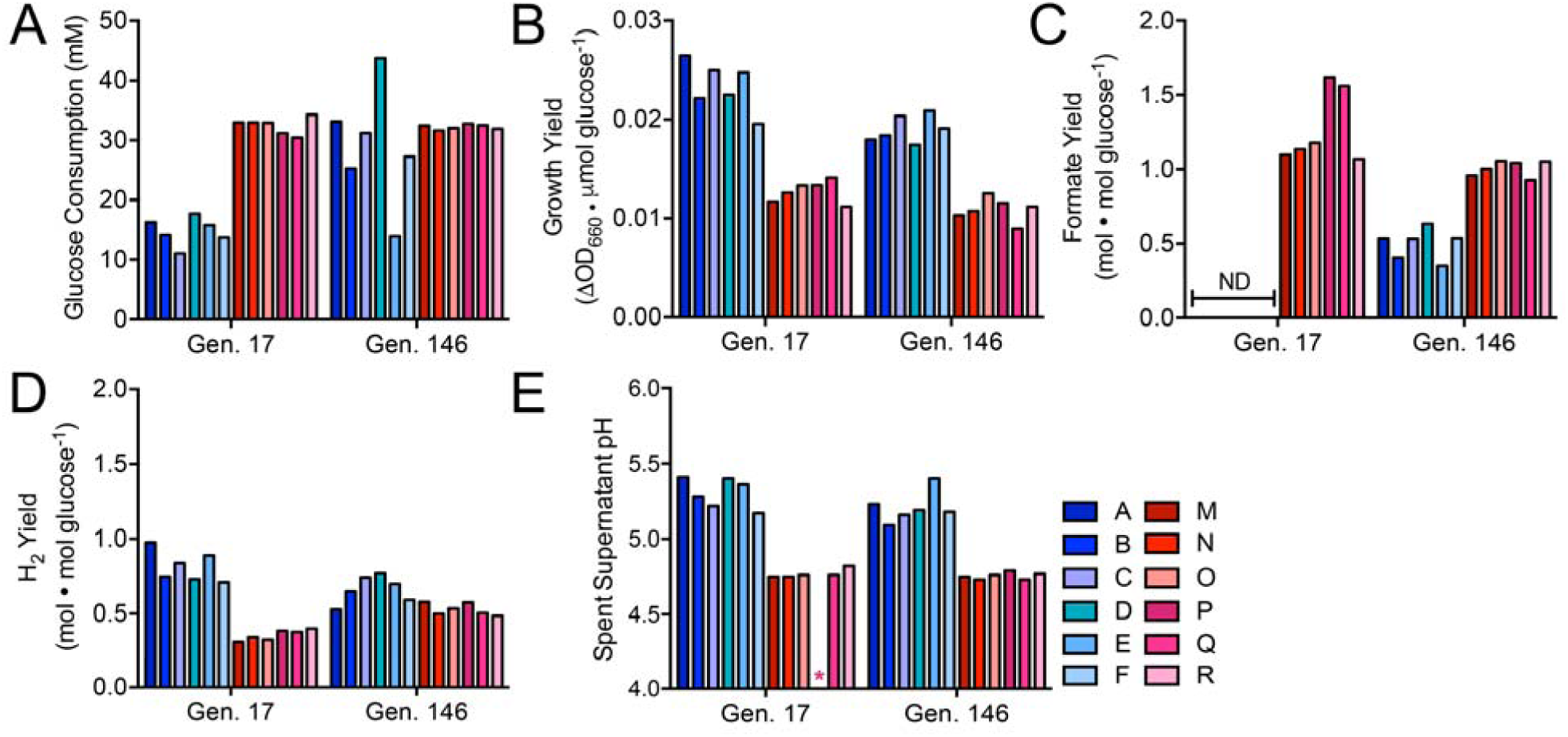
WT-based (blue) and NifA*-based (red) cocultures exhibit distinct metabolic phenotypes. Bars represent a single measurement for each lineage for glucose consumption (A), growth yield (B), formate yield (C), H_2_ yield (D), and final pH (E) for the indicated WT-based (CGA4001; blue) and NifA*-based (CGA4003; red) revived coculture lineages at generation (Gen) 17 and 146. Different shades indicate different lineages. ND, not detected. Asterisk (*) indicates that the pH for lineage P at G17 was not quantified because culture tube broke prior to measurement.

We observed a potential trade-off between coculture growth rate and coculture growth yield (ΔOD_660_/glucose consumed). For example, WT-based G17 cocultures had the slowest growth rates but highest growth yields, whereas NifA*-based G146 cocultures had the fastest growth rates but lowest growth yields (Fig. 4B). Trade-offs between growth rate and yield have been reported in multiple microbial species under various conditions [39-41]. In our case, the metabolic trends point to possible explanations for the apparent trade-off. For example, formate produced by *E. coli* is not consumed by *R. palustris* and thus typically accumulates in cocultures [17]. However, no formate was detected in WT-based G17 cocultures and formate yields were approximately half that of NifA*-based cocultures at G146 (Fig. 4C). Low formate yields could be explained in part by increased conversion of formate to H_2_ and CO_2_ by *E. coli* formate hydrogenlyase [42, 43]. Consistent with this possibility, WT-based cocultures had the highest H_2_ yields (Fig. 4D). Low formate yields could also be explained by decreased formate production by *E. coli* in favor of other fermentation products. We previously observed low formate yields in slow-growing, nitrogen-limited NifA*-based cocultures [19, 20], suggesting that formate production by *E. coli* varies in response to growth rate. We have also not ruled out the possibility that *R. palustris* can consume some formate under certain conditions. In addition to formate, consumable organic acid yields were also lower at both G17 and G146 for WT-based cocultures relative to NifA*-based cocultures (Fig. S2). Organic acid accumulation in cocultures can acidify the medium to inhibitory levels [17]. At both G17 and G146, the lower yields of formate and other organic acids in WT-based cocultures translated into higher pH values than in NifA*-based cocultures (Fig. 4E). This lower level of acidification combined with the likelihood of a higher proportion of glucose being fermented into organic acids other than formate could explain the higher *R. palustris* cell densities at G146 in WT-based cocultures compared to NifA*-based cocultures (Fig. 3B).

### A single mutation in an *E. coli* nitrogen starvation response regulator is sufficient for mutualistic growth with WT *R. palustris*

We hypothesized that the growth of WT-based cocultures was due to adaptive mutations in one or both species. To determine whether the evolution of either or both species was necessary to establish a nascent mutualism, we isolated single colonies of each species from ancestral WT populations and evolved G146 cocultures and paired them in all possible combinations (Fig. 5A). Only those pairings featuring evolved *E. coli* grew to an OD_660_ >0.5 after ∼24 days (Fig. 5B). Cocultures pairing evolved *E. coli* with ancestral or evolved WT *R. palustris* exhibited similar doubling times of ∼67 h (Fig. S1). These results indicate that adaptation by *E. coli* alone is sufficient to establish a nascent mutualism with WT *R. palustris*. Accordingly, we did not observe increased NH_4_^+^ excretion in evolved WT *R. palustris* N_2_-fixing monocultures compared to the ancestral strain (Fig. S1).

**Fig. 5.**
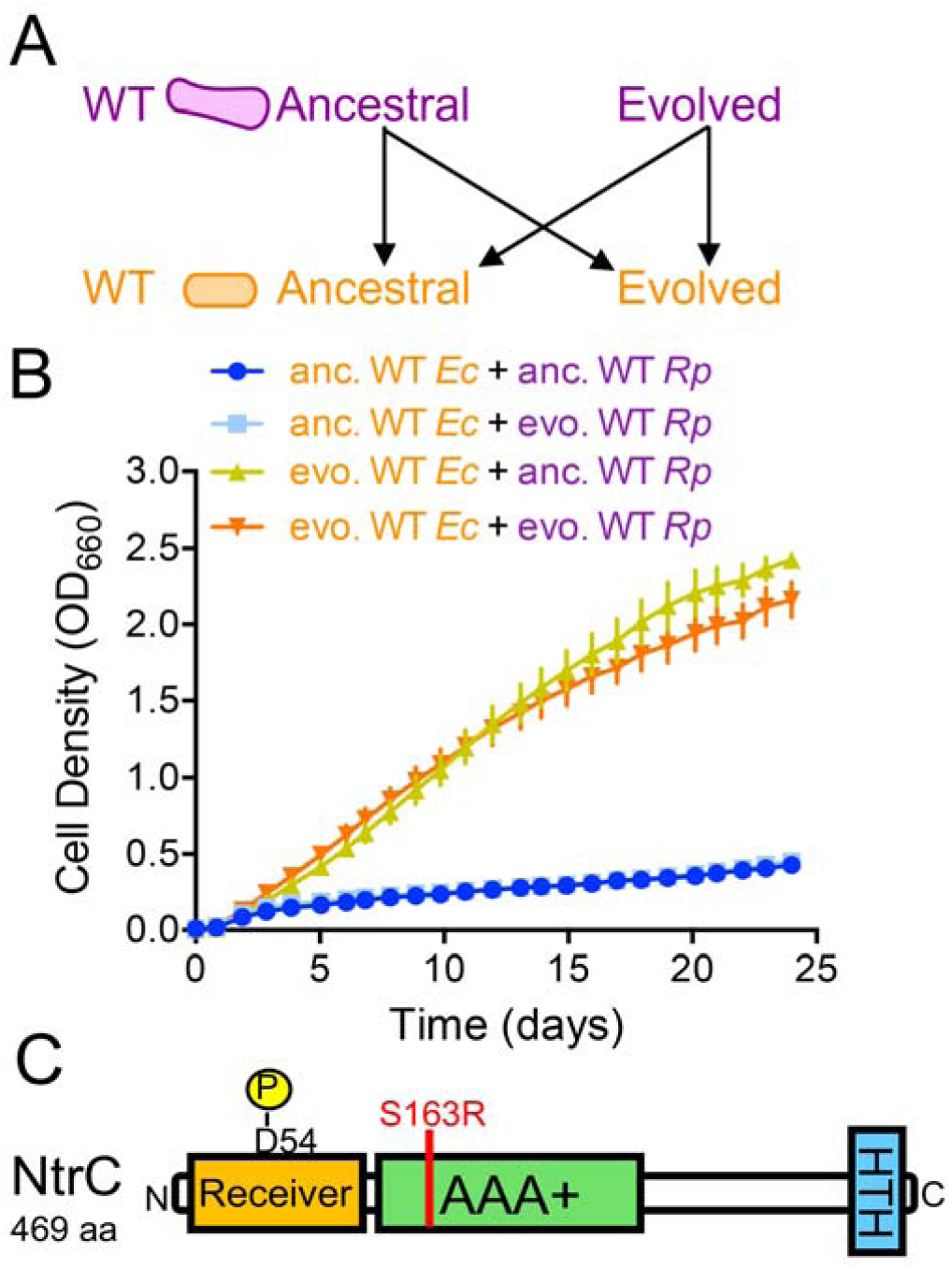
Adaptation by *E. coli* is sufficient to enable growth of WT-based cocultures. Ancestral (anc) and evolved (evo) WT *R. palustris* (CGA4001) and WT *E. coli* were paired in all possible combinations (A) and the growth of the cocultures (both species) was monitored (B). (B) Points are means ± SEM, n=3. (C) The location (red line) of the missense mutation in *E. coli* NtrC, which was fixed in all six parallel evolved *E. coli* populations from WT-based cocultures at G140-146.

To identify candidate mutations in *E. coli* that could drive improved coculture growth, we sequenced the genomes of populations in each evolved coculture lineage after 123-146 generations. We also sequenced WT-based cocultures following ∼11 generations to determine if potentially adaptive mutations arose early within WT-based cocultures. Several parallel mutations were identified in both species at frequencies between 5 and 100% (Table 1 and Supplementary Files 1 and 2). Consistent with evolved *E. coli* being necessary for mutualistic growth with either ancestral or evolved WT *R. palustris* (Fig. 5B), we did not detect any *nifA* nor *amtB* mutations in evolved WT *R. palustris* populations, which would enable rapid coculture growth. The multiple high frequency parallel mutations observed in evolved *R. palustris* populations (Table 1), are insufficient to improve coculture growth trends (Fig. 5B). Of the mutations in evolved *E. coli* populations, we were intrigued by a fixed missense mutation in *glnG* (henceforth called *ntrC*) that occurred in all evolved *E. coli* populations from both shaken and static WT-based cocultures, replacing serine 163 with an arginine within the AAA+ domain in the encoded the response regulator NtrC (NtrC^S163R^, Table 1 and Fig. 5C). NtrC and the histidine kinase NtrB form a two-component system that senses and coordinates the nitrogen starvation response in *E. coli* [44-46]. Our lab previously found that the *E. coli* NtrBC-regulon is highly expressed in coculture with *R. palustris* NifA* [20]. Thus, *E. coli* NtrBC might be even more important in coculture with WT *R. palustris* wherein *E. coli* nitrogen starvation is expected to be intensified.

**Table 1.**
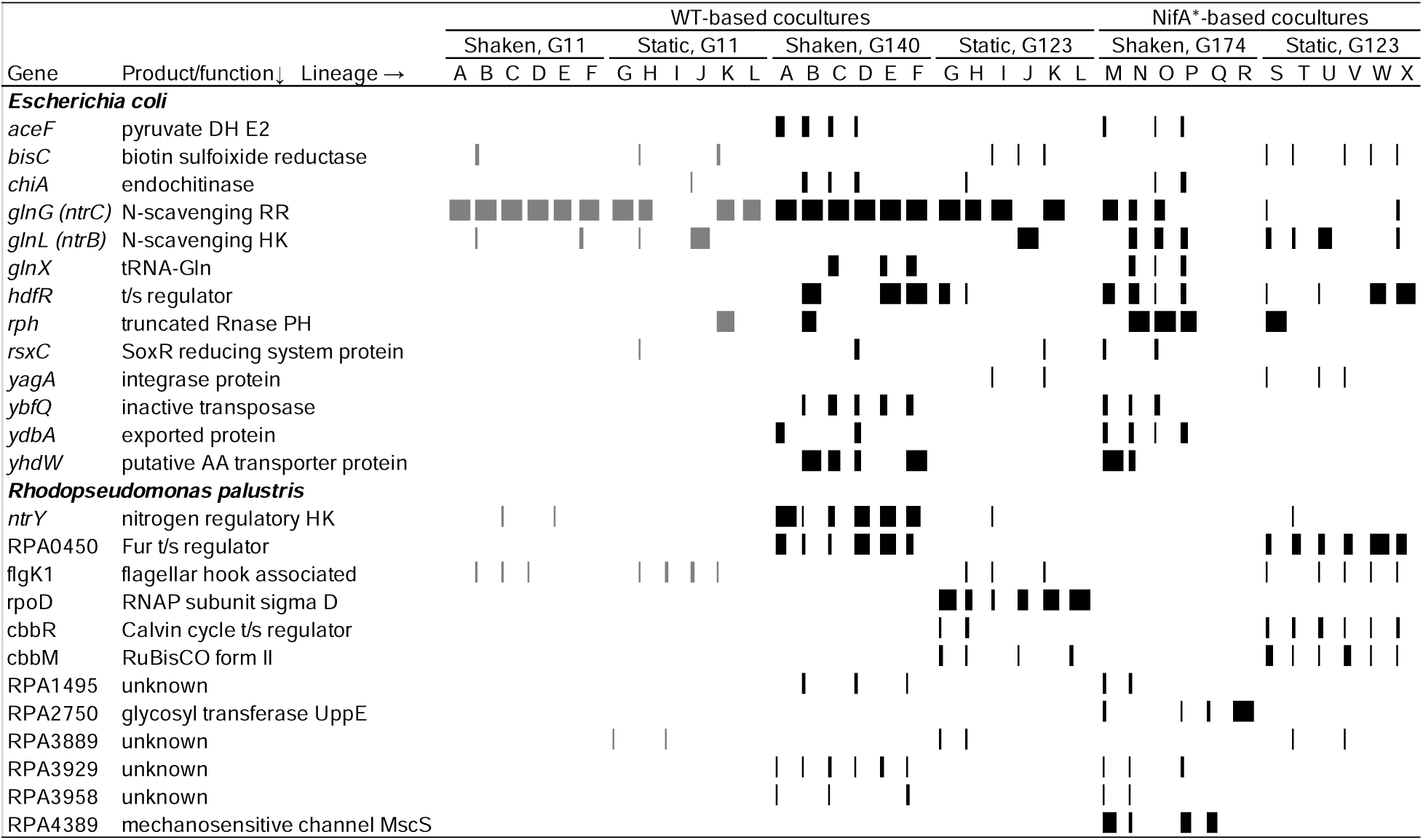
Parallel mutations in evolved WT-based and NifA*-based cocultures. Genes included have > 1 missense mutation, premature stop, insertion, and/or deletion in > 4 lineages, have passing scores, and occur at frequency of > 5% within a population at the later time point (black bars). Bar width corresponds to mutation frequency. Frequencies at the early time point for the genes with the above criteria are also shown (gray bars). The highest frequency was used when multiple mutations were identified in a single gene within a single lineage. Coverage was too low to detect mutations in E. coli lineages Q and R. Details of these and other mutations, including intergenic mutations with high parallelism that were omitted here for simplicity, are in Supplementary Files 1 and 2. DH, dehydrogenase; HK, histidine kinase; RR, response regulator; t/s transcription.

The NtrC^S163R^ mutation was enriched early in the evolution of WT-based cocultures, already at a high frequency in most lineages by G11 (Table 1 and Supplementary Table 3). Because of the striking parallelism of the NtrC^S163R^ mutation across coculture lineages, we wondered if it was present as standing genetic variation in the ancestral *E. coli* population. We therefore sequenced *ntrC* of ten *E. coli* isolates subjected to a single round (35 days) of coculture growth with WT *R. palustris* (Fig. 6A), which should enrich for the NtrC^S163R^ mutation. All ten isolates sequenced had the WT *ntrC* allele. Thus, although the NtrC^S163R^ allele was likely present in the founder population given its presence in every WT-based coculture lineage, it was likely under strong selection from an initial low frequency. In support of the importance of the NtrC^S163R^ allele, we also identified multiple, though different, high frequency mutations in *ntrB* and *ntrC* in *E. coli* populations from NifA*-based cocultures evolved under both well-mixed and static conditions (Table 1 and Supplementary Table 3). Together, these observations strongly suggest the adaptive importance of *E. coli ntrBC* mutations like NtrC^S163R^ for coculture growth, regardless of the NH_4_^+^-excreting phenotype of the *R. palustris* partner.

**Fig. 6.**
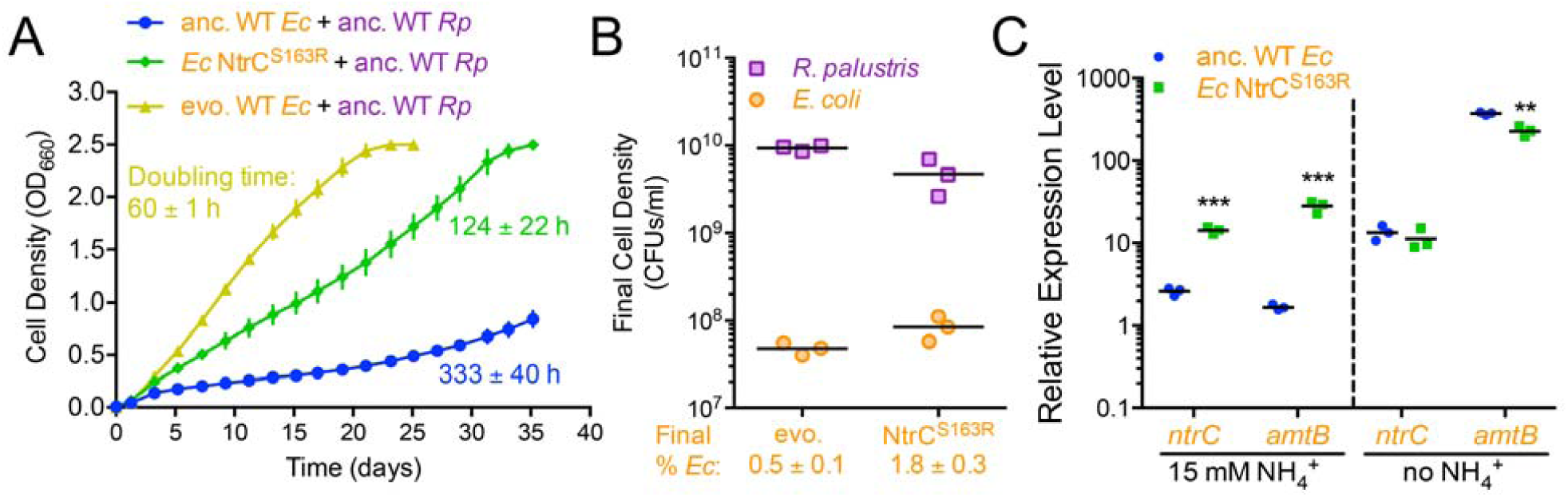
A missense mutation in *E. coli ntrC* enables emergent NH_4_^+^ cross-feeding by conferring constitutive expression of nitrogen acquisition. (A) Coculture growth curves (both species) of ancestral (anc) WT, evolved (evo) WT, and the NtrC^S163^ mutant *E. coli* paired with ancestral WT *R. palustris* (CGA4001). Points are means ± SEM, n=3. Mean doubling times (± SD) are listed next to each growth curve. (B) Final cell densities of each species and *E. coli* frequencies in cocultures with evolved WT *E. coli* and the NtrC^S163^ mutant at the final time points shown in panel A. Triplicate technical replicate plating was performed for each biological replicate. Final *E. coli* frequencies are the mean ± SD. (C) Relative expression of *ntrC* and *amtB* genes in ancestral WT *E. coli* and the NtrC^S163^ mutant when grown in monoculture with 15 mM NH_4_Cl or under complete NH_4_Cl starvation. (B, C) Points represent biological replicates and lines are means, n=3; Holm-Sidak t-test, \*\**p*<0.01, \*\*\**p*<0.001). RT-qPCR experiments were performed with duplicate technical replicates for each biological replicate. *E. coli hcaT* was used for normalization. Similar results were observed with *gyrB* and with multiple primer sets for both the target and reference housekeeping genes.

To determine if the NtrC^S163R^ mutation alone was sufficient to support coculture growth with WT *R. palustris*, we moved the NtrC^S163R^ allele into the ancestral *E. coli* strain. Cocultures pairing *E. coli* NtrC^S163R^ with WT *R. palustris* grew with a doubling time of 124 ± 22 h, approximately twice as long as cocultures with evolved *E. coli* isolates, but much faster than cocultures with ancestral *E. coli* (Fig. 6A). Thus, the NtrC^S163R^ mutation is sufficient to drive coculture growth. Cocultures with *E. coli* NtrC^S163R^ reached similar final cell densities, but supported lower WT *R. palustris* abundances than cocultures with evolved *E. coli* isolates (Fig. 6B).

We speculate that some of the additional parallel mutations in evolved *E. coli* (Table 1) are also adaptive and account for the faster coculture growth rate relative to cocultures with the NtrC^S163R^ mutant (Fig. 6A). For example, the *E. coli* strain we used has a frameshift mutation in *rph* that decreases expression of *pyrE* immediately downstream [47]. Mutations in *rph* and in-between *rph* and *pyrE*, like those identified here (Table 1 and Supplementary Files 1 and 2), can improve pyrimidine biosynthesis and growth in minimal media [48]. HdfR is a transcriptional regulator that inhibits flagellar expression [49] and activates glutamate synthase expression [50]. HdfR loss-of-function mutations could reduce glutamate synthase expression and synchronize NH_4_^+^ assimilation with the low NH_4_^+^ cross-feeding levels in WT-based cocultures. However, the entire HdfR regulon has never been reported and thus could include other genes. In another possible link to nitrogen metabolism, mutations accumulated in the glutamine tRNA gene, *glnX* (Table 1). Two of the mutations disrupt the base-pair adjacent to the anti-codon loop while two others alter the anti-codon, one still coding for glutamine but the other coding for histidine. The impact of these mutations is difficult to predict, especially since GlnX is just one of four *E. coli* tRNAs for glutamine. The *yhdWXYZ* operon encodes an NtrC-regulated amino acid ABC transporter, which is predicted to be non-functional due to a frameshift mutation in *yhdW* [44, 45]. The mutations identified in (Table 1) and directly upstream of the *yhdW* pseudogene (Supplementary Files 1 and 2) could restore amino acid transport for use as a nitrogen source or alternatively disrupt NtrC binding upstream of *yhdW* and thereby free up NtrC to regulate genes more critical to NH_4_^+^ acquisition.

### The *E. coli* NtrC^S163R^ allele constitutively activates ammonium transporter expression

Based on the effects of NtrC mutations observed by others [51, 52], we hypothesized that the NtrC^S163R^ allele facilitates coculture growth with WT *R. palustris* by conferring constitutive expression of NtrBC-regulated genes important for NH_4_^+^ acquisition. We previously determined that NtrC and AmtB were crucial gene products within the NtrBC regulon for growth and coexistence with *R. palustris* NifA* [18, 20]. To test if the NtrC^S163R^ allele increased *amtB* and *ntrC* expression, we measured transcript levels by reverse transcription quantitative PCR (RT-qPCR) in *E. coli* monocultures grown with 15 mM NH_4_Cl or subjected to complete nitrogen starvation (∼10 h with 0 mM NH_4_Cl). We chose to perform RT-qPCR on *E. coli* monocultures, because ancestral WT *E. coli* does not readily grow with WT *R. palustris* and because *E. coli* typically constitutes a low percentage (1-5%) of WT-based cocultures, meaning most mRNA would be from *R. palustris*.

When cultured with NH_4_Cl, the *E. coli* NtrC^S163R^ mutant exhibited ∼30 and ∼15-fold higher expression of *amtB* and *ntrC*, respectively, than WT *E. coli* (Fig. 6C), indicating that the NtrC^S163R^ allele constitutively activates expression of its regulon. Following 10 h of nitrogen starvation, we saw similarly high *amtB* and *ntrC* expression by both the WT and the NtrC^S163R^ strains (Fig. 6C). Thus, both the WT and NtrC^S163R^ *E. coli* strains are able to commence strong transcriptional responses to extreme nitrogen starvation. We expect that the level of nitrogen limitation experienced by *E. coli* in coculture with WT *R. palustris* is less extreme than the complete nitrogen starvation conditions used in our qPCR experiments. Although we cannot detect NH_4_^+^ excretion by WT *R. palustris*, the equilibrium with NH_3_ dictates that some will be excreted, possibly within the nM to low µM range where AmtB is critical [21]. We also know that AmtB is important in coculture for *E. coli* to compete for transiently available NH_4_^+^ that *R. palustris* will otherwise reacquire [18]. We therefore hypothesize that the NtrC^S163R^ mutation primes *E. coli* for coculture growth with *R. palustris* by maintaining high AmtB expression and thereby enabling acquisition of scarcely available NH_4_^+^. The resulting faster *E. coli* growth and metabolism would translate into faster organic acid excretion, which itself would foster *R. palustris* growth and reciprocal NH_4_^+^ excretion. Thus, we envision that better NH_4_^+^ acquisition creates a positive feedback loop, enhancing the growth of both players in the mutualism.

## DISCUSSION

Here, we determined that in cocultures requiring nitrogen transfer from *R. palustris to E. coli*, an *E. coli* NtrC^S163R^ mutation alone is sufficient to enable coculture growth. The mutation results in constitutive activity of the NtrC regulon and thus increased expression of the AmtB NH_4_^+^ transporter, which we hypothesize enhances NH_4_^+^ uptake. This is the first mutation we have identified in the NH_4_^+^ recipient *E. coli* that is sufficient to support mutualistic growth with WT *R. palustris*. Overall, our data suggest that a recipient species can persuade cross-feeding through enhanced nutrient uptake. An improvement in nutrient acquisition by one species could lead to increased reciprocation and thus generate a positive feedback loop in the context of mutualism.

Our previous work on this consortium utilized *R. palustris* NifA* and ΔAmtB strains that we engineered to excrete NH_4_^+^ [17-19]. In the present study, we did not identify *nifA* or *amtB* mutations in evolved WT *R. palustris* populations. *R. palustris nifA* and *amtB* mutations likely incur a fitness cost, such as an increased energetic burden of constitutive nitrogenase expression due to the NifA* mutation or loss of NH_4_^+^ to WT competitors in the case of an inactivating *amtB* mutation. Thus, emergent *R. palustris nifA* and *amtB* mutants would not be expected to be competitive in the presence of a large WT *R. palustris* population. However, it does not appear that *R. palustris* NifA* regained regulation of nitrogenase and limited NH_4_^+^ excretion during the experimental evolution of NifA*-based cocultures. Instead, ancestral and evolved NifA*-based cocultures supported consistent abundances of *E. coli*, a trait that is dependent on the level of NH_4_^+^ excretion [17-19]. Our results therefore suggest that the NifA* mutation, a 48-bp deletion, is not prone to frequent or rapid suppression, at least during the time scale of this study, potentially because multiple mutations would be required. It is also possible that the cost of constitutive N_2_ fixation is relatively low within this mutualism. Based on our findings, we propose that experimental evolution of synthetic consortia is useful for both for identifying novel genotypes enabling coexistence and for assessing the stability of putatively costly engineered genotypes.

More broadly, our results indicate that within a cross-feeding partnership, multiple combinations of recipient and producer genotypes can lead to stable coexistence but only certain combinations will be favored based on the selective environment. Under well-mixed conditions there is intense competition between recipients as well as producers for limiting, communally-valuable nutrients, such as NH_4_^+^ [18], vitamins, or amino acids [1, 7, 8]. Additionally, there is a probable fitness cost for producers associated with increased nutrient excretion in well-mixed environments. Under well-mixed conditions, costless self-serving and mutually beneficial mutations, but not costly partner-serving mutations, are favored to evolve [53]. Therefore mutations that improve a recipient’s ability to acquire nutrients from producers, and thereby outcompete other recipient genotypes, can evolve rapidly [54]. Recipient mutations that enhance metabolite uptake also erode the partial privatization of communally valuable nutrients released by the producer [55]. Even so, these recipient mutations can benefit producers if they promote mutualistic interactions [54]. We view the *E. coli* NtrC^S163R^ mutation as an example of a conditionally-costless self-serving mutation, given its rapid emergence in well-mixed cocultures, but one that is mutually beneficial in the context of an obligate mutualism. We hypothesize that the benefit of the NtrC^S163R^ mutation for *E. coli* extends more generally to surviving nitrogen limitation. In support of this, an NtrC^V18L^ mutation that similarly increased *amtB* expression was adaptive for *E. coli* evolved in nitrogen-limiting monocultures [56]. *E. coli* is nitrogen-limited in all coculture conditions used in this study, likely explaining why *E. coli* NtrBC mutations were also observed in evolved NifA*-based cocultures, and in all static cocultures, where the dense populations at the bottom of the test tube likely intensify competition for NH_4_^+^.

Mutations that improve nutrient acquisition can be mutually beneficial for cross-feeding partners under conditions where neither species can grow well without reciprocal nutrient exchange. However, mutations that enhance nutrient uptake could also be adaptive for the recipient when there is no reciprocal benefit to the producer. For example, acetate cross-feeding repeatedly evolved in *E. coli* populations under glucose-limiting conditions through mutations that enhanced acetate uptake by a nascent recipient subpopulation [57, 58]. Unlike in our study, the acetate-consuming recipients did not provide a clear reciprocal benefit to the acetate-excreting producers beyond a potentially relaxed competition for glucose due to resource partitioning [57, 58]. Consequently, mutations that enhance nutrient uptake could foster the emergence of mutualistic, commensal, or competitive interactions, depending on community composition and conditions [54, 57, 58]. In natural microbial communities, where auxotrophy is prevalent [1, 7] and most cells exhibit low metabolic activity [9, 10], mutations that improve acquisition of limiting nutrients could allow certain populations to flourish. Understanding the consequences of mutations that expedite metabolite acquisition could thus inform on the origins of various ecological relationships. This knowledge could ultimately be harnessed for applications ranging from facilitating coexistence within synthetic consortia to probiotic-mediated competitive exclusion of pathogens.

## Acknowledgements

This work was supported in part by U.S. Army Research Office grants W911NF-14-1-0411 and W911NF-17-1-0159, a National Science Foundation CAREER award MCB-1749489, the U.S. Department of Energy, Office of Science, Office of Biological and Environmental Research, under award DE-SC0008131, and the Joint Genome Institute Community Science Program, CSP 502893. The work conducted by the U.S. Department of Energy Joint Genome Institute, a DOE Office of Science User Facility, is supported by the Office of Science of the U.S. Department of Energy under Contract No. DE-AC02-05CH11231.

We thank AL Posto, JR Gliessman, and MC Onyeziri for coculture passaging and initial characterizations, JT Lennon and BK Lehmkuhl for equipment and assistance with qRT-PCR, and J Ford and AM Buechlein at the IU Center for Genomics and Bioinformatics.

## SUPPLEMENTARY INFORMATION

**Supplemental File 1. Mutations identified in evolved WT-based cocultures A25, B24-F24 and NifA*-based cocultures M30-R30 using *breseq*.**

**Supplemental File 2. Mutations identified in evolved WT-based cocultures A1-F1 (mixed/shaking), G1-L1 (static), G21-L21 (static), and NifA*-based cocultures S21-X21 (static) using the BBMap package**

**Fig. S1.**
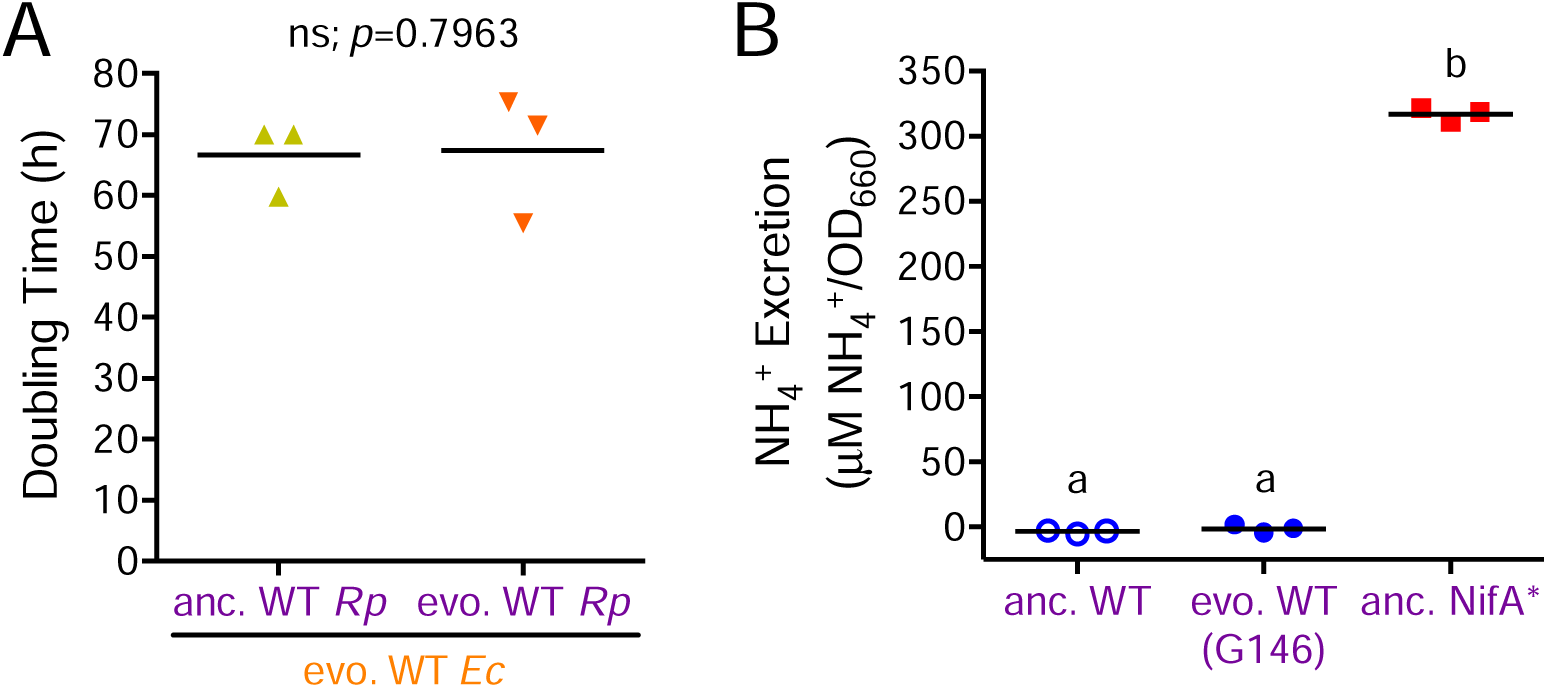
Evolution of WT *R. palustris* in coculture with *E. coli* does not affect coculture doubling times nor NH_4_^+^ excretion levels. (A) Coculture doubling time of evolved WT *E. coli* (G146, A25 isolates) paired with ancestral or evolved WT *R. palustris* (CGA4001; G146, A25 isolates). Points represent biological replicates and lines are means, n=3; paired t-test, *p*=0.7963; ns, not significant). (B) NH_4_^+^ excretion by ancestral and evolved WT *R. palustris* and the NifA* mutant during carbon-limited N_2_-fixing monoculture growth. Points represent biological replicates and lines are means, n=3; One-way ANOVA with Tukey’s multiple comparisons test, different letters indicate significant statistical differences, p<0.0001).

**Fig. S2.**
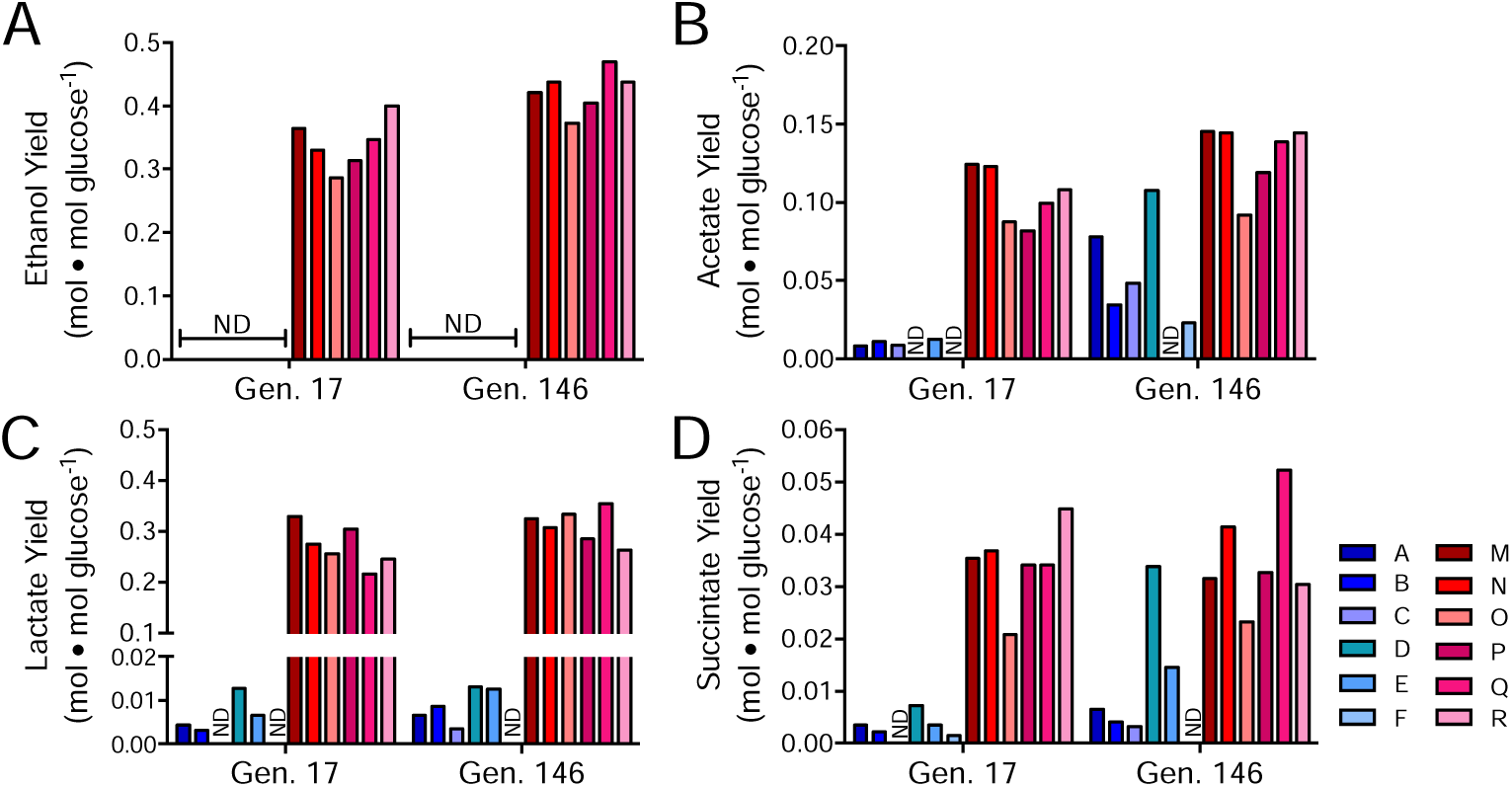
Other evolved coculture fermentation product yields also differ between WT-based and NifA*-based cocultures. Bars are individual yields for ethanol (A), acetate (B), lactate (C), and succinate (D), for the indicated WT-based (CGA4001; blue) and NifA*-based (CGA4003; red) revived coculture lineages at generation (Gen) 11 and 146. Different shades indicate different lineages. ND, not detected.

**Supplementary Table S1.** Strains and plasmids.

| Strain or plasmid | Genotype (text designation); Phenotype/description | Reference, origin, or description |
| --- | --- | --- |
| *R. palustris* strains |  |  |
| CGA009 | Wild-type strain; spontaneous Cm<sup>R</sup> derivative of CGA001 | [59] |
| CGA676 | NifA* (NifA*); derivative of CGA009 with 48 bp deletion of NifA Q-linker (amino acids 202-217) conferring constitutive nitrogenase expression | [24] |
| CGA4001 | $\Delta hupS$ (anc. WT); Ancestral strain for experimentally evolved WT-based cocultures A-F; derivative of CGA009 with inactive uptake hydrogenase | This study |
| CGA4003 | *nifA\** $\Delta hupS$ (anc. NifA*); Ancestral strain for experimentally evolved NifA*-based cocultures M-R; derivative of CGA009 with constitutive nitrogenase expression and inactive uptake hydrogenase | This study |
| *E. coli* strains |  |  |
| MG1655 | Wild-type K-12 strain (WT/anc.); ancestral *E. coli* strain for all experimental evolution lineages | [60] |
| MG1655 NtrC<sup>S163R</sup> | NtrC<sup>S163R</sup> (NtrC<sup>S163R</sup>); MG1655 derivative with serine 163 to arginine (S163R) point mutation of NtrC | This study |
| S17-1 | *thi pro hdsR hdsM<sup>+</sup> recA*; chromosomal insertion of RP4-2 (Tc::Mu Km::Tn7); | [61] |
| Plasmids |  |  |
| pJQ200SK |  | [27] |
| pJQ-$\Delta hupS$ | pJQ200SK with DNA fragments flanking *hupS* fused by PCR to generate unmarked, in-frame deletion of *hupS* in *R. palustris* | [17] |
| pKD46 | Cb<sup>R</sup>; temperature-sensitive plasmid with arabinose-inducible λ-Red recombination system for recombineering of *E. coli* | [28] |

**Supplementary Table S2.**
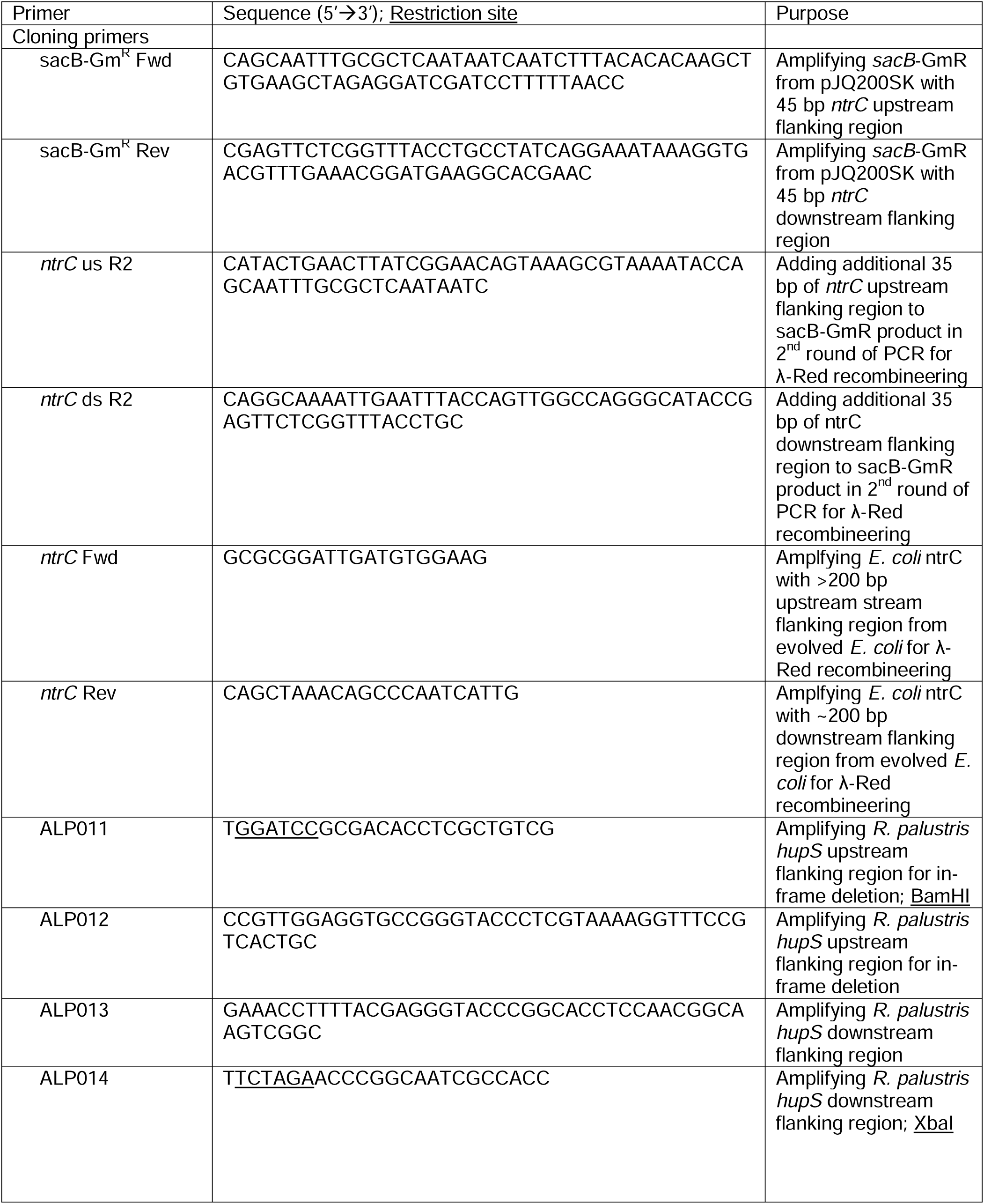
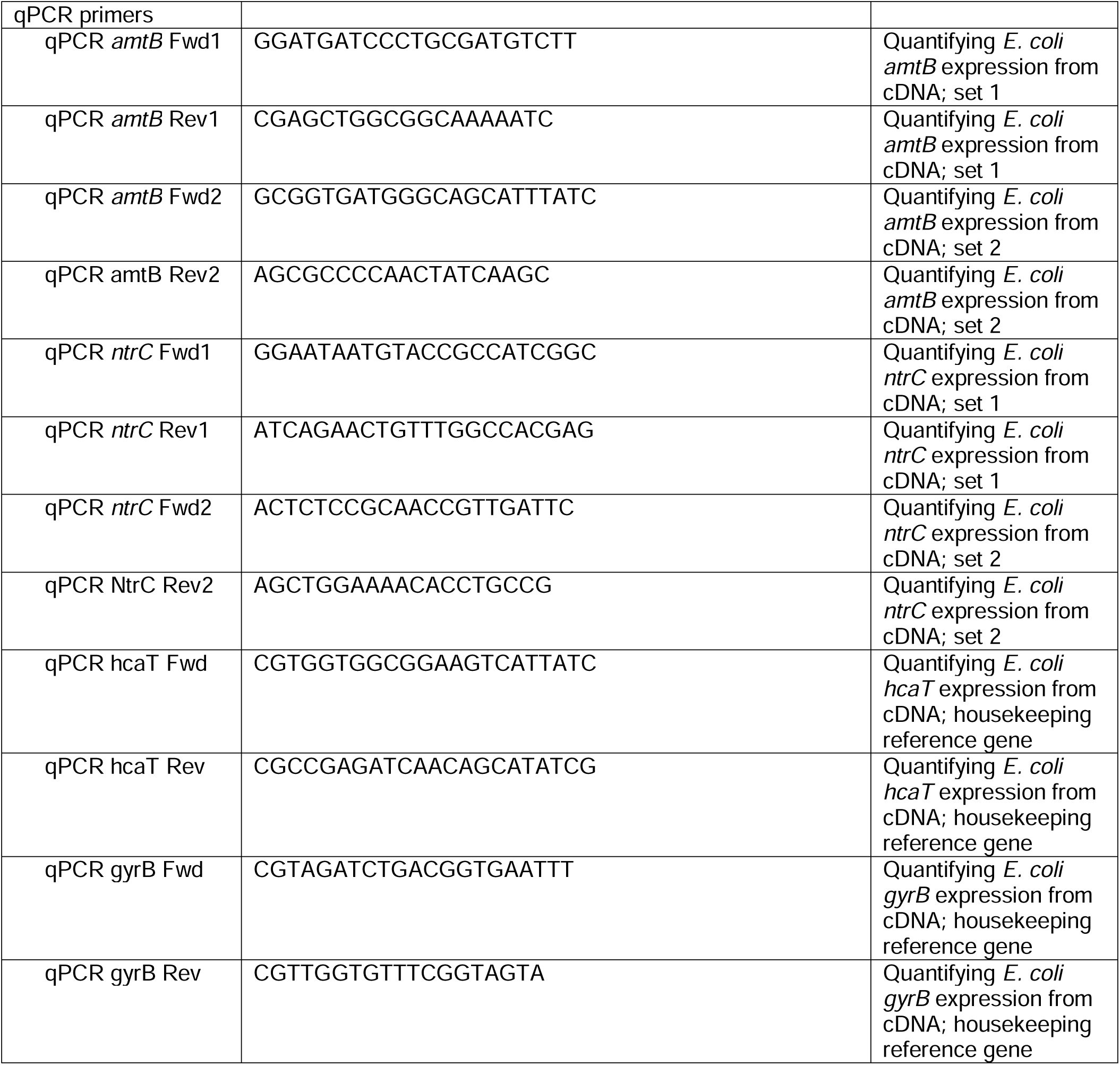
Primers.

**Supplementary Table 3.** Mutations in *ntrBC* genes in *E. coli* following coculture evolution.

| Lineage | Gene | Mutation | Generation | Frequency | *R. palustris*<br>partner strain | Growth<br>condition | SRA<br>Accession |
| --- | --- | --- | --- | --- | --- | --- | --- |
| A | *ntrC* | S163R | 11 | 100% | WT | Mixed | SRX5772396 |
| A | *ntrC* | S163R | 146 | 100% | WT | Mixed | SRX5872514 |
| B | *ntrC* | S163R | 11 | NC<sup>a</sup> | WT | Mixed | SRX5772395 |
| B | *ntrC* | S163R | 140 | 100% | WT | Mixed | SRX5872520 |
| C | *ntrC* | S163R | 11 | NC<sup>a</sup> | WT | Mixed | SRX5772261 |
| C | *ntrC* | S163R | 140 | 100% | WT | Mixed | SRX5874533 |
| D | *ntrC* | S163R | 11 | NC<sup>a</sup> | WT | Mixed | SRX5772258 |
| D | *ntrC* | S163R | 140 | 100% | WT | Mixed | SRX5874537 |
| E | *ntrC* | S163R | 11 | NC<sup>a</sup> | WT | Mixed | SRX5772266 |
| E | *ntrC* | S163R | 140 | 100% | WT | Mixed | SRX5874556 |
| F | *ntrC* | S163R | 11 | NC<sup>a</sup> | WT | Mixed | SRX5209606 |
| F | *ntrC* | S163R | 140 | 100% | WT | Mixed | SRX5874560 |
| G | *ntrC* | S163R | 11 | NC<sup>a</sup> | WT | Static | SRX5772260 |
| G | *ntrC* | P399T | 11 | 17% | WT | Static | SRX5772260 |
| G | *ntrC* | S163R | 123 | 100% | WT | Static | SRX5772001 |
| H | *ntrC* | S163R | 11 | 63% | WT | Static | SRX5772264 |
| H | *ntrC* | S163R | 123 | 76% | WT | Static | SRX5771991 |
| I | *ntrC* | S163R | 11 | NC<sup>a</sup> | WT | Static | SRX5772259 |
| I | *ntrC* | S163R | 123 | 100% | WT | Static | SRX5771995 |
| J | *ntrB* | R116S | 11 | NC<sup>a</sup> | WT | Static | SRX5772262 |
| J | *ntrB* | R116S | 123 | 100% | WT | Static | SRX5771992 |
| K | *ntrC* | S163R | 11 | 80% | WT | Static | SRX5772263 |
| K | *ntrC* | S163R | 123 | 100% | WT | Static | SRX5771996 |
| L | *ntrC* | S163 | 11 | NC<sup>a</sup> | WT | Static | SRX5772265 |
| L | ND<sup>b</sup> | - | 123 | - | WT | Static | SRX5209608 |
| M | *ntrC* | H184Y | 174 | 79% | NifA* | Mixed | SRX5874674 |
| N | *ntrB* | A175V | 174 | 39% | NifA* | Mixed | SRX5875647 |
| N | *ntrC* | D109V | 174 | 10% | NifA* | Mixed | SRX5875647 |
| N | *ntrC* | G373V | 174 | 44% | NifA* | Mixed | SRX5875647 |
| O | *ntrB* | P336Q | 174 | 41% | NifA* | Mixed | SRX5877494 |
| O | *ntrC* | Q374H | 174 | 54% | NifA* | Mixed | SRX5877494 |
| P | *ntrB* | R116H | 174 | 32% | NifA* | Mixed | SRX5877497 |
| Q | ND<sup>b</sup> | - | 174 | - | NifA* | Mixed | SRX5910505 |
| R | ND<sup>b</sup> | - | 174 | - | NifA* | Mixed | SRX5910551 |
| S | *ntrB* | L137V | 123 | 23% | NifA* | Static | SRX5771990 |
| S | *ntrC* | S45R | 123 | 7% | NifA* | Static | SRX5771990 |
| T | *ntrB* | G150C | 123 | 15% | NifA* | Static | SRX5771989 |
| U | *ntrB* | A175V | 123 | 65% | NifA* | Static | SRX5771988 |
| V | ND<sup>b</sup> | - | - | - | NifA* | Static | SRX5771994 |
| W | ND<sup>b</sup> | - | - | - | NifA* | Static | SRX5771993 |
| X | *ntrB* | A175V | 123 | 13% | NifA* | Static | SRX5771967 |
| X | *ntrB* | V305F | 123 | 16% | NifA* | Static | SRX5771967 |
| X | *ntrC* | E116K | 123 | 17% | NifA* | Static | SRX5771967 |

<sup>a</sup>Frequency was not calculated (NC) because read coverage <10.<sup>b</sup>*ntrBC* mutations were not detected (ND) in population. Mutations present at frequencies <1% are not listed.

